# ConoDL: A Deep Learning Framework for Rapid Generation and Prediction of Conotoxins

**DOI:** 10.1101/2024.09.27.614001

**Authors:** Menghan Guo, Zengpeng Li, Xuejin Deng, Ding Luo, Jingyi Yang, Yingjun Chen, Weiwei Xue

**Author notes:** **Corresponding Authors** Dr. Weiwei Xue, Dr. Yingjun Chen. M.G. and Z.L. contributed equally to this work.

## Abstract

Conotoxins, being small disulfide-rich and bioactive peptides, manifest notable pharmacological potential and find extensive applications. However, the exploration of conotoxins’ vast molecular space using traditional methods is severely limited, necessitating the urgent need of developing novel approaches. Recently, deep learning (DL)-based methods have advanced to the molecular generation of proteins and peptides. Nevertheless, the limited data and the intricate structure of conotoxins constrain the application of deep learning models in the generation of conotoxins. We propose ConoDL, a framework for the generation and prediction of conotoxins, comprising the end-to-end conotoxin generation model (ConoGen) and the conotoxin prediction model (ConoPred). ConoGen employs transfer learning and a large language model (LLM) to tackle the challenges in conotoxin generation. Meanwhile, ConoPred filters artificial conotoxins generated by ConoGen, narrowing down the scope for subsequent research. A comprehensive evaluation of the peptide properties at both sequence and structure levels indicates that the artificial conotoxins generated by ConoDL exhibit a certain degree of similarity to natural conotoxins. Furthermore, ConoDL has generated artificial conotoxins with novel cysteine scaffolds. Therefore, ConoDL may uncover new cysteine scaffolds and conotoxin molecules, facilitating further exploration of the molecular space of conotoxins and the discovery of pharmacologically active variants. The code is available at https://github.com/xueww/ConoDL.The supplementary material and model are available at https://zenodo.org/records/10679280.

## Introduction

Conotoxins, rich-cysteine peptides derived from the cone snail venom, typically comprise 10 to 70 amino acid residues and feature multiple pairs of disulfide bonds^1^. Conotoxins exhibit significant pharmacological potential, as they can interact with multiple targets, such as voltage-gated ion channels, ligand-gated ion channels, G-protein-coupled receptors, and neurotransmitter transports^2^. Currently, Ziconotide (ω-conotoxin MVIIA) has obtained clinical approval for the treatment of chronic pain^3^, and several other conotoxin-based drugs are in various stages of clinical trials^4^. It is estimated that there are approximately 10^6^ conotoxins in existence, but less than 1% of them have been described to date, and even fewer have been pharmacologically characterized^5–7^. Existing methods for exploring conotoxins include purification and characterization from venom of cone snails^8, 9^, chemical modification or structural alteration of existing conotoxin^10, 11^, and total synthesis of conotoxins^12, 13^. Although conotoxins occupy a vast molecular space, existing methods for exploring the spatial aspects of conotoxins are highly limited. Therefore, novel approaches to facilitate the discovery of new conotoxins are urgently needed.

Recently, as artificial intelligence (AI) technology develops, especially the advanced deep learning (DL)-based methods including ProteinGAN^14^, ProGen^15^, ProtGPT^16^, ProteinMPNN, etc., have achieved significant success in the molecular generation of proteins and peptides^17^. ProteinGAN expands the protein molecular space through the application of generative adversarial networks (GANs)^14^. ProGen rapidly generates protein sequences spanning multiple families with large language model (LLM)^15^. ProtGPT generates protein in a high-throughput manner using a self-regressive model based on the generative pretrained transformer (GPT) architecture^16^. ProteinMPNN de novo design protein sequences employing message passing neural network (MPNN)^18^. In addition, models based on the Wasserstein autoencoder (WAE) can capture semantic relationships between peptide sequences, leading to a significant acceleration in the discovery functional peptides such as antimicrobial peptides (AMPs)^19^ and anticancer peptides (ACPs)^20^. Therefore, multiple DL algorithms present great potential to develop powerful tools to facilitate the survey of the conotoxins’ vast molecular space.

It is remarkable that the development of DL methods in the molecular generation of conotoxins faces certain challenges as follows: (1) The number of known conotoxins may be insufficient for DL training, with existing data comprising only around two thousand sequences^21^; (2) Conotoxin sequences exhibit a wide range of lengths, with the shortest being only 6 amino acids and the longest exceeding a hundred amino acids^22^; and (3) The structural diversity of conotoxins poses a challenge, with more than 30 disulfide bond scaffolds in conotoxins, making it difficult to learn internal patterns within the molecular space^1, 23^. However, transfer learning reduces the reliance on data and is particularly well-suited for datasets with limited data and complex intrinsic patterns, such as the conotoxins dataset^24^. The pre-training LLMs using the universal protein sequences such as ProGen has captured the fundamental intrinsic patterns of proteins^15^ and holds the potential to tackle the challenges associated with applying DL to the molecular generation of conotoxins. Fine-tuning of the pre-training model with conotoxins dataset may enable a quicker and more effective acquisition of the intricate intrinsic patterns of conotoxins.

In this study, we propose a framework ConoDL for the rapid and large-scale generation and preliminary identification of conotoxins. ConoDL consists of two modules: ConoGen and ConoPred. Firstly, a transformer-based conotoxins generation model ConoGen, was trained using mature peptides collected from ConoServer^22^. ConoGen can rapidly and efficiently produce artificial conotoxins. Subsequently, a prediction model ConoPred based on the WAE architecture^20^, was trained to predict the probability that the generated sequences are conotoxins. By establishing a probability threshold, sequences that are less likely to be authentic conotoxins can be eliminated. Finally, we conducted a comprehensive comparison between artificial conotoxins and natural conotoxins in terms of both sequence and spatial aspects. At the sequence level, artificial conotoxins exhibit similarities with natural conotoxins in terms of sequence length, cysteine framework, amino acid distribution, charge distribution, and hydrophobicity distribution. At the spatial level, the cysteine residues in artificial conotoxins are capable of forming disulfide bonds, leading to the adoption of specific spatial conformations. These evaluation results demonstrate the feasibility and superiority of ConoDL in exploring the conotoxin landscape. Therefore, ConoDL has the potential to reveal previously undiscovered conotoxin molecules and new cysteine scaffolds, thus advancing further exploration of the molecular space of conotoxins and aiding in the subsequent discovery of pharmacologically active variants.

## Results and discussion

### Pipeline of conotoxins generation, selection, and *in silico* evaluation

The comprehensive pipeline is illustrated in **Figure 1**, providing an overview of our approach. Initially, we utilized the Conotoxin dataset along with large language model (LLM) ProGen^15^ through transfer learning techniques to train ConoGen, a conotoxin generation model engineered for rapid and customized synthesis of conotoxins. Subsequent to this, we employed both the conotoxin dataset and additional protein dataset to train a conotoxin prediction model, ConoPred, which was applied to predict the artificial conotoxins generated by ConoGen. ConoGen and ConoPred collectively constitute the deep learning framework, ConoDL, for conotoxin generation and prediction. Finally, a comprehensive evaluation was conducted to assess the artificial conotoxins. At the sequence level, various aspects were considered, including sequence length distribution, cysteine residue distribution, amino acid distribution, physicochemical properties, and similarity. At the structural level, structural prediction tools were firstly employed to construct and compare the folding types of sequences generated by ConoDL. Then, all atom molecular dynamics (MD) simulations were utilized to assess the structural stability of these sequences. The multi-level analysis from sequence to structure lays the groundwork for further research and optimization.

**Figure 1.**
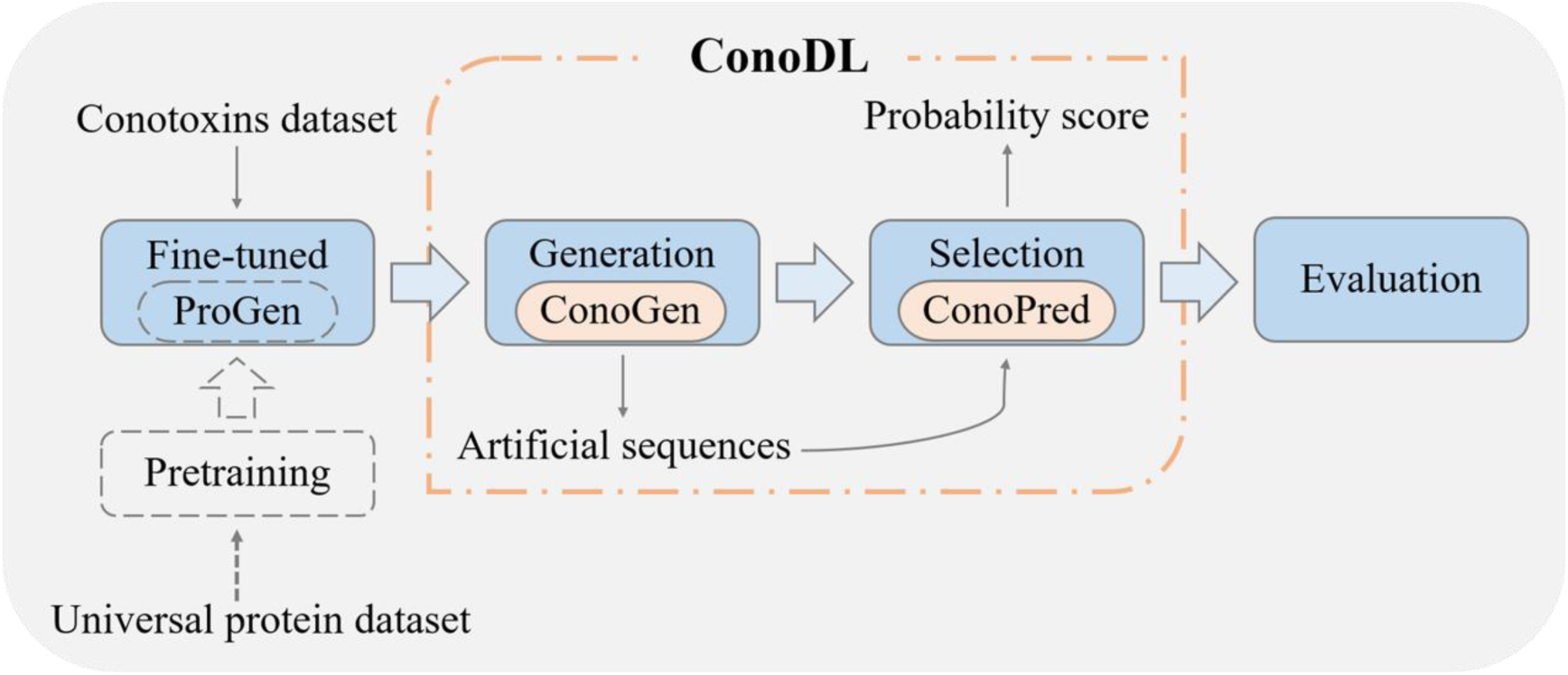
Schema of the computational modules in this study. ConoDL, formed by ConoGen and ConoPred. ConoGen was first obtained by fine-tuning the pretraining large language model (LLM) ProGen^15^ using the sequences from Conotoxin dataset. Then, the generated sequences of conotoxins from ConoGen were subjected to preliminary filtering by the trained prediction model ConoPred. And in silico methods were used to further evaluate the sequence and structure profiles of potential conotoxins.

### Framework of ConoDL

As described in the pipeline (Figure 1), ConoDL consists of two modules: (1) ConoGen, a conotoxin generation model for generating artificial conotoxin sequences; and (2) ConoPred, a conotoxin prediction model that outputs the probability score of whether the input sequence is a conotoxin. The following is the architecture details of the ConoGen and ConoPred in our ConoDL.

#### Architecture of ConoGen

ConoGen is a conotoxin generation model that adopts an architecture consistent with the pre-training model ProGen (Figure 2A)^15^. ConoGen is also constructed using a Transformer-based neural network architecture. One major advantage of the Transformer lies in its self-attention mechanism, which enables the model to encode distant signals of information when making sequence predictions, thereby affording it the capability to comprehend complex semantics. Its Transformer architecture consists of 36 layers, each comprising 8 self-attention heads, totaling 1.2 billion trainable parameters to ensure the training and optimization of the model.

**Figure 2.**
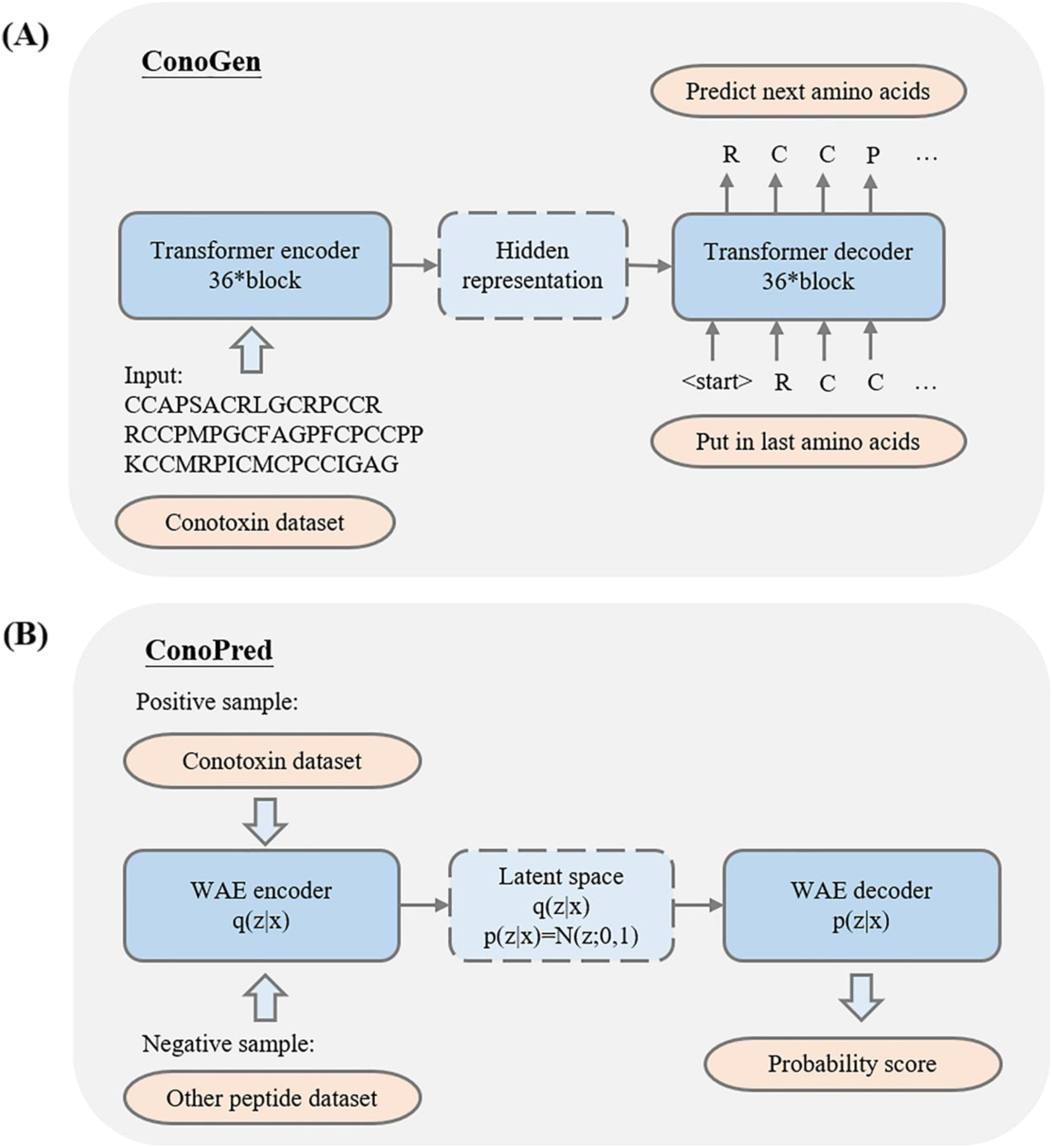
Framework of ConoDL. **(A)** The architecture diagram of the ConoGen model trained in this work. ConoGen is a neural network based on the Transformer architecture, which use a self-attention mechanism to model interactions among amino acid residues. ConoGen generates artificial conotoxin sequences by predicting the probability distribution of the next amino acid and minimizing the loss in the next amino acid prediction task. **(B)** The architecture diagram of the ConoPred model trained in this work. ConoPred is a neural network based on the Wasserstein Autoencoder (WAE) architecture. The encoding process involves mapping input data q(x|z) to the latent space z of the model and constructing a density model for the latent space p(z│x)=N(z;0,1). The decoding process reconstructs the original data based on the learned distribution.

ConoGen provides the probability distribution *p*(*x*) for predicting the next amino acid based on context^25^:

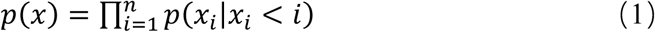

Then generates artificial conotoxin sequences by minimizing the negative log-likelihood *L*(*B*) associated with the next amino acid prediction task^26^.

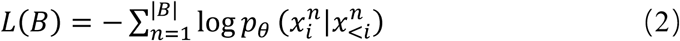

During the generation process, various hyperparameters can be configured to control the diversity, length, and cysteine scaffold of artificial conotoxin sequences by altering the probability distribution of predicting the next amino acid.

#### Architecture of ConoPred

ConoPred is a conotoxin prediction model that adopts an architecture based on Wasserstein AutoEncoder (WAE) (Figure 2B). The goal of WAE is to train a model that can map input data to a latent space (encoding) and reconstruct the original data from it (decoding). Simultaneously, it minimizes the Wasserstein distance to ensure similarity between the distribution in the latent space and a predefined prior distribution. The Wasserstein distance is a method for measuring the difference between two distributions. In contrast to traditional autoencoders, WAE differs by introducing a prior on the latent distribution and employing the Wasserstein distance to measure the distance between the encoded distribution and the prior distribution. The Wasserstein distance is expressed as follows:

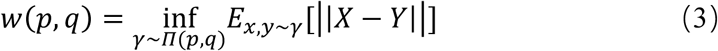

*Π* represents the set of all possible joint distributions combining the distributions *p* and *q*; ||*X* − *Y*|| denotes the distance between samples *X* and *Y* sampled from the joint distribution. The Wasserstein distance is the lower bound on the expected value of this distance.

To address the issue of class imbalance, we employed the Focal Loss function in combination with the Adam optimizer to update the model parameters. The Focal Loss function is expressed as follows:

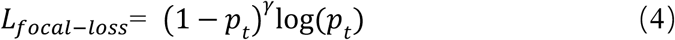

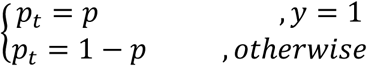

*p_t_* represents the probability predicted by the model for the input to be a positive sample, and *γ* is the balance coefficient used to adjust the balance between positive and negative samples.

### Generation ability of ConoGen

ConoGen is a generation model designed for the rapid generation of artificial conotoxin sequences. This model can generate ten thousand artificial conotoxin sequences within ten minutes. Even for tasks with relatively limited training data, ConoGen can effectively leverage the available data for learning. In our work, we employed ConoGen to rapidly generate a batch of 1 million artificial conotoxin sequences. After removing duplicates, we obtained 761,371 non-redundant sequences, resulting in a redundancy rate of only 23.9%.

### Prediction ability of ConoPred

ConoPred is a prediction model trained based on the WAE architecture. When a sequence is fed into ConoPred, it generates a probability score that estimates the probability of that sequence being classified as a conotoxin. The ConoPred was employed to predict the probability scores for 439 conotoxin sequences from the ConoServer database that were not included in the training set, 11,531 other sequences from the UniProt database that also excluded from training, as well as 761,371 artificial conotoxin sequences genera ted by ConoGen. The corresponding probability scores for all the sequences were shown in Figure 3. It is observed that the probability scores for conotoxins are predominantly above 0.90, while the probability scores for other sequences mostly remain below 0.10. To some extent, this attests to the accuracy of the prediction model ConoPred. Meanwhile, the probability scores for the artificial conotoxin sequences generated by ConoGen largely exceeded 0.80, thereby indicating that the artificial conotoxin sequences generated by ConoGen conform to the internal patterns of conotoxins.

**Figure 3.**
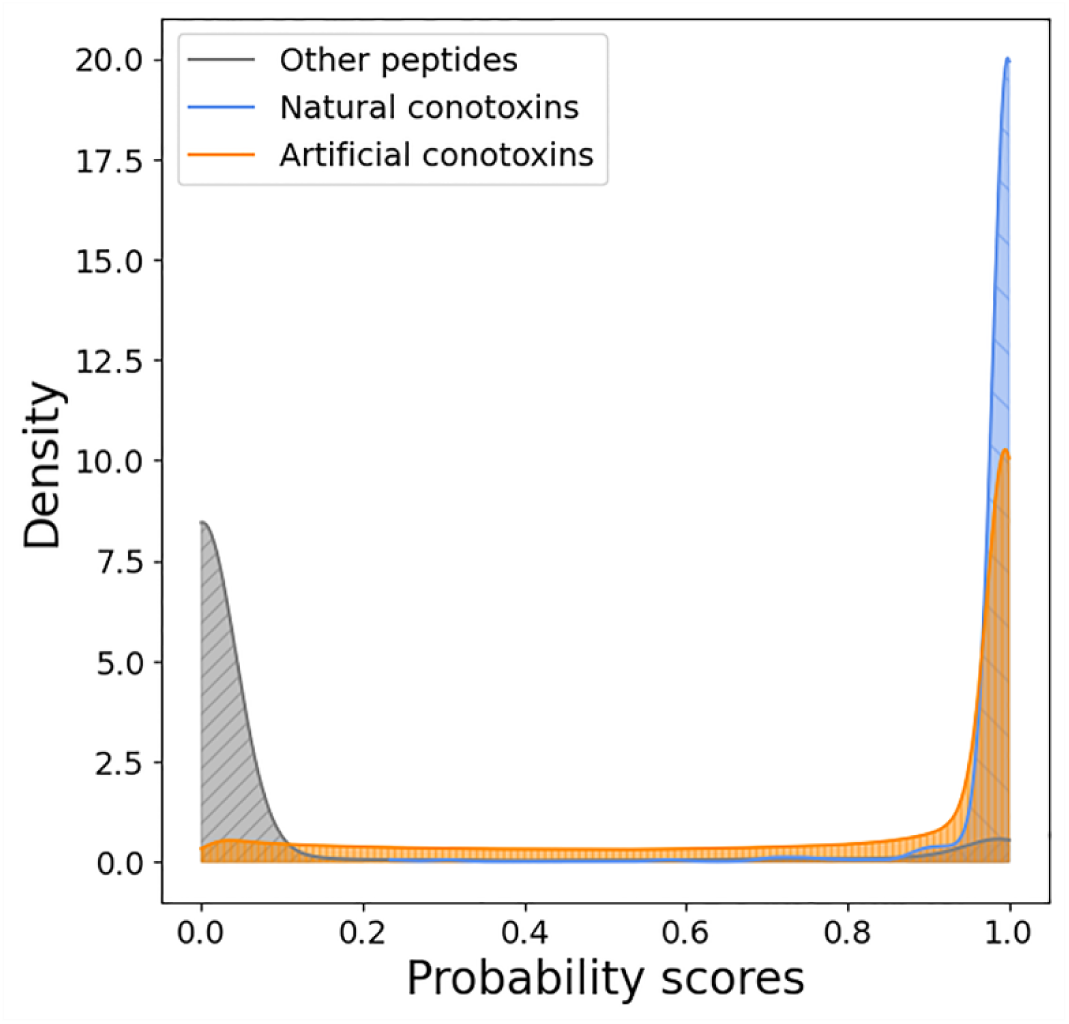
Prediction ability of ConoPred reflected by probability scores on three different datasets. Plot of sequence density versus conotoxin probability score (gray: Other sequences in the UniProt database (non-training set); blue: Conotoxins from the ConoServer database (non-training set); orange: Artificial conotoxin sequences generated by ConoGen).

### Comprehensive evaluation of artificial and natural conotoxin sequences

From sequence perspective, we conducted a comprehensive analysis of both artificial and natural conotoxin sequences, including the length distribution, cysteine scaffold types, physicochemical properties, and the similarity.

#### Sequence length

The length distribution of artificial peptides generated by ConoGen was analyzed and depicted in Figure 4A. The majority (80%) of peptides fall within the length ranges from 20 to 60. The length distribution range of artificial peptides generated by ConoGen is generally consistent with that in the ConoServer database (Figure 4B). However, ConoGen’s ability to generate shorter artificial conotoxin sequences (10-20 amino acids) is lacking. This may be related to the following factors: (1) Complexity of the model. Complex models typically have larger memory capacity, allowing them to capture longer-term dependencies^27^. The complexity of the ConoGen leads to a tendency for generating longer sequences as it can maintain more internal information. 2) Complexity of the data. If the training data includes sequences of various lengths, the model learns to capture this diversity and endeavors to reflect this characteristic during the generation process^28^. The sequences in the conotoxin dataset vary in length, and ConoGen is designed to generate sequences of different lengths to accommodate this diversity. 3) Randomness in the generation process. ConoGen uses the softmax function for sampling when generating sequences, enabling the selection of the next amino acid. This randomness contributes to variations in the length of the generated peptide sequences within a certain range^29^. The above description indicates that the majority of artificial conotoxins generated by ConoGen align with the sequence length range of natural conotoxins, demonstrating that ConoGen has captured the intrinsic patterns of natural conotoxins in terms of sequence lengths.

**Figure 4.**
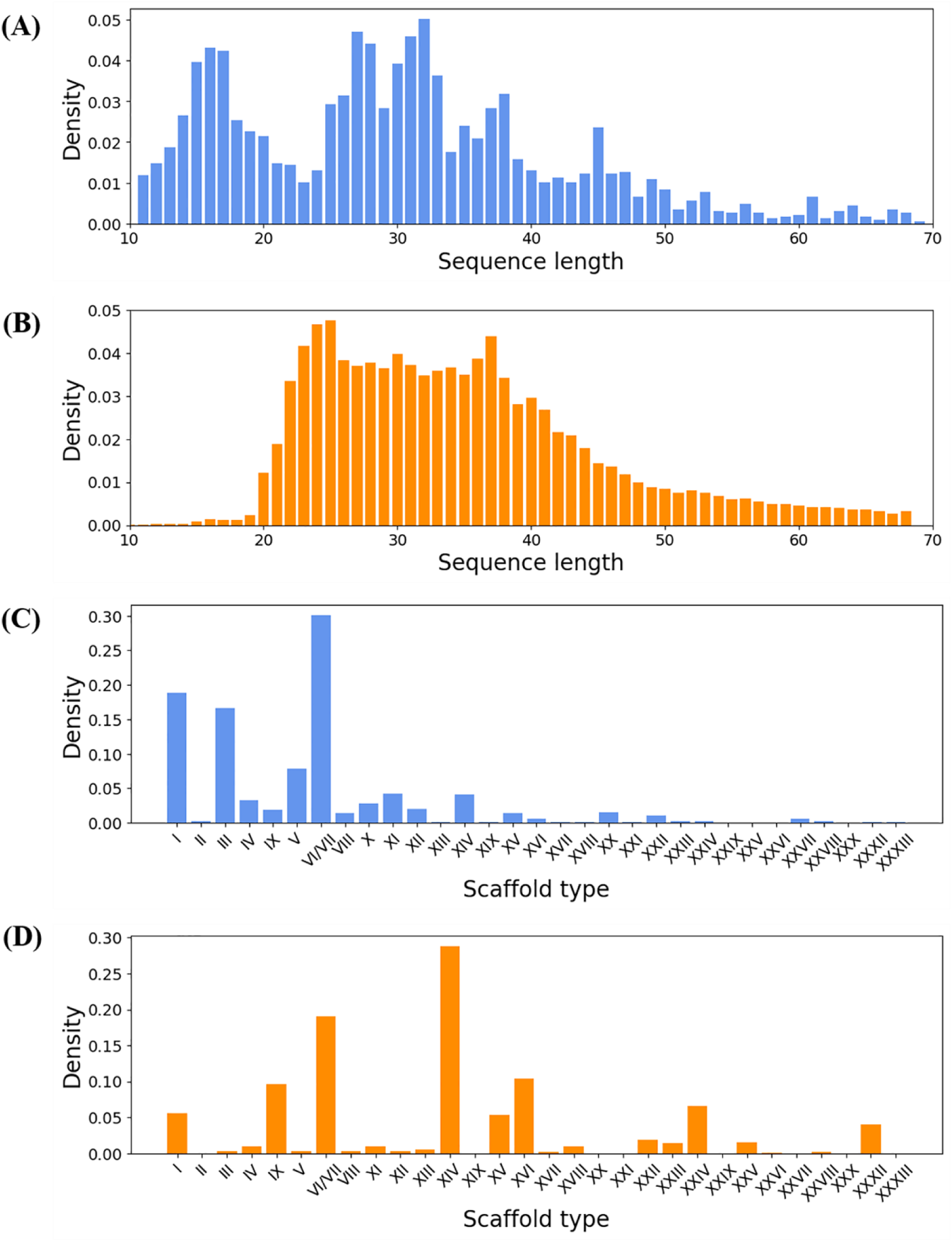
Comparison of sequence length and cysteine framework types between artificial and natural conotoxins. **(A)** Length distribution of mature peptide sequences in the ConoServer database. **(B)** Length distribution of artificial conotoxin sequences generated by ConoGen. **(C)** Distribution of cysteine scaffold types in mature conotoxin peptide sequences within the ConoServer database. **(D)** Distribution of cysteine scaffold types in artificial conotoxin sequences generated by ConoGen.

#### Cysteine scaffold

A total of 31 natural occurring cysteine frameworks have been identified from the ConoServer database (Figure 4C). Analyzing the distribution of cysteine scaffold types shown that although two cysteine frameworks, namely X (CC-C.[PO]C) and XXIX (CCC-C-CC-C-C), was not generated under the current set of hyperparameters, approximately 35.8% of the generated conotoxin sequences conform to the 29 existing cysteine scaffold types of conotoxins (Figure 4D), while the remaining 64.2% artificial sequences may contain undiscovered new scaffold types. Remarkably, the number of sequences for 8 framework types exceed 5,000, while for 18 framework types exceed 1,000. Thus, in addition to the existing cysteine scaffolds, the artificial conotoxins generated by ConoGen involve specific scaffolds, with the potential for the development of artificial conotoxins with novel structures.

#### Physicochemical properties

For the sequences of artificial conotoxin generated by ConoGen and conotoxins from the ConoServer database, the physicochemical properties (Figure 5) including amino acid composition, charge distribution, and hydrophobicity were calculated and compared. In (Figure 5A), the 373490 artificial conotoxin sequences showed a similar distribution of amino acids with the natural conotoxin sequences. However, compared with the natural conotoxins, the amino acid distribution was not entirely identical. The sequences generated by ConoGen were rich in Cysteine (Cys), and they exhibited slightly higher levels of Glycine (Gly) and Serine (Ser), along with lower levels of Methionine (Met) and Tryptophan (Trp), indicating a greater sequence diversity of possibilities. Furthermore, artificial conotoxin sequences generated by ConoGen matched well with conotoxins in terms of charge distribution (Figure 5B) and hydrophobicity distribution (Figure 5C). It indicates that ConoGen can successfully capture the inherent patterns of physicochemical properties in conotoxins.

**Figure 5.**
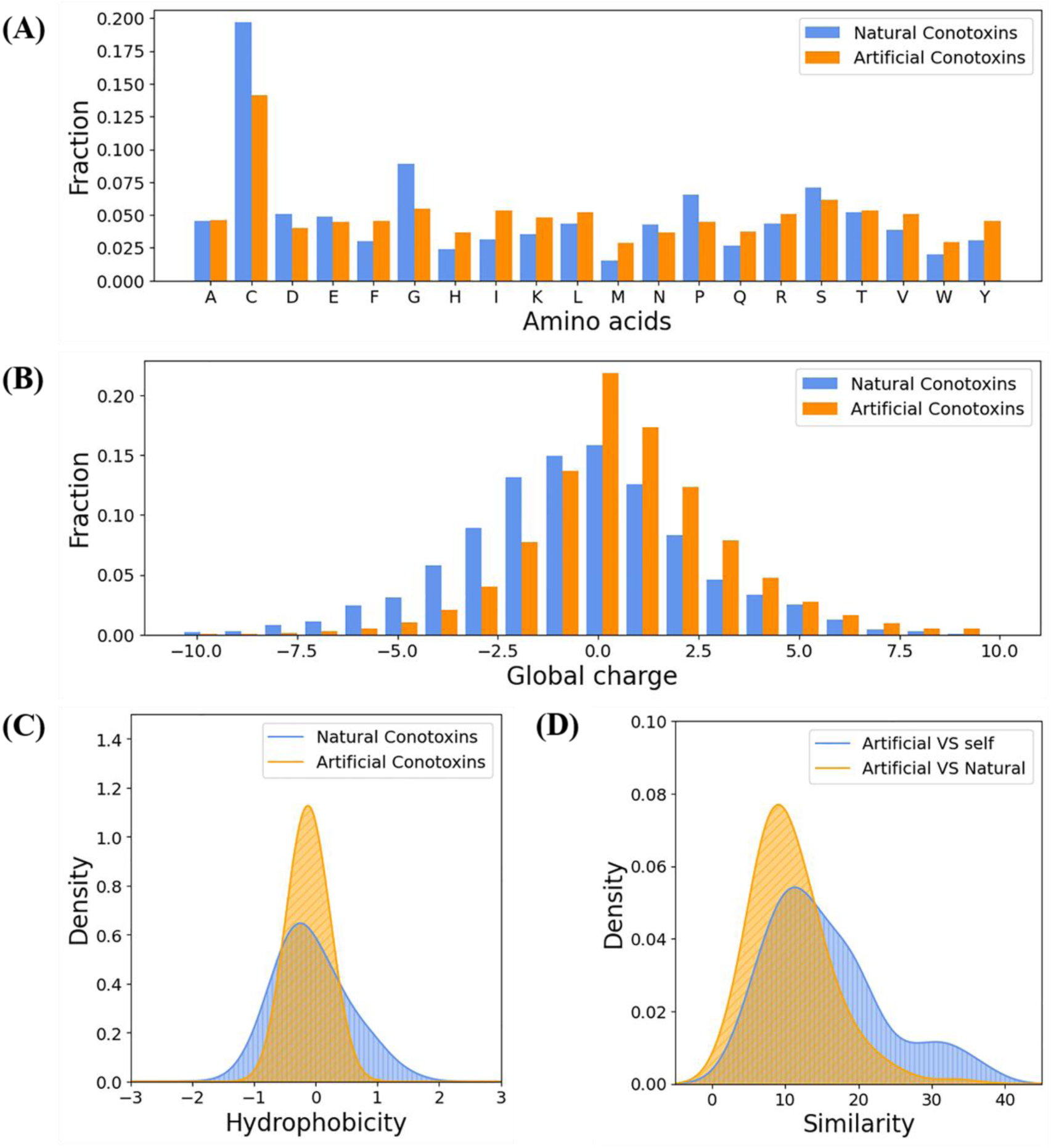
Comparison of amino acid composition, charge distribution, hydrophobicity, and sequence similarity between artificial and natural conotoxins. **(A)** The amino acid distributions (blue: conotoxins from the ConoServer database; orange: artificial conotoxin sequences generated by ConoGen). **(B)** The net charge distributions (blue: Conotoxins from the ConoServer database; orange: Artificial conotoxin sequences generated by ConoGen). **(C)** The hydrophobicity distribution (blue: Conotoxins from the ConoServer database; orange: Artificial conotoxin sequences generated by ConoGen). **(D)** The sequence similarity between conotoxins (orange: comparison between generated sequences and the dataset; blue: self-comparison of generated sequences).

#### Sequence similarity

The sequence similarity is crucial for generative models as it serves to evaluate the learning capability of the model and the innovativeness of the sequences it generates. From a statistical perspective, when employing the globalds function to assess two entirely identical sequences, the average similarity score is 36.8. In contrast, for two completely dissimilar sequences, the globalds function provides an average similarity score of −14.6^20^. The similarity between the training set (conotoxins in the ConoServer database) and sequences generated by ConoGen, as well as the similarity of ConoGen-generated sequences self, is shown in Figure 5D. According to the statistics, the average similarity between the generated sequences and the training set is 10.86, while the average similarity of the generated sequences self is 15.58. It is reasonable to infer that ConoGen has captured the internal features among conotoxins, and concurrently generated sequences also exhibit a degree of novelty, making them effective tools for exploring conotoxins.

### 3D structures and dynamic profiles of the artificial conotoxins

From a spatial perspective, the formation of disulfide bonds in different artificial conotoxins was first analyzed, and with the conformational changes were further investigated by relatively long time-scale molecular dynamics (MD) simulations at atomic-levels.

#### Specific disulfide bonds in the 3D structures of different artificial conotoxins

The oxidation of cysteine residues leading to the formation of disulfide bonds is crucial for maintaining the conformation of proteins. Through this conformation, disulfide bonds play a pivotal role in the stability and activity of proteins. Consequently, whether cysteine residues in artificial conotoxins can form disulfide bonds and fold into 3D structures similar to those of natural conotoxins constitutes a crucial aspect in evaluating the generation quality of ConoGen^30^. Taking conotoxins with different cysteine scaffold types as examples, we constructed their 3D structures. The 3D structures of ConoGen-generated artificial conotoxins and those of natural conotoxins were compared and shown in **Table 1**. It is observed that the artificial conotoxins generated by ConoGen not only exhibit specific cysteine scaffolds at the sequence level, but also form disulfide bonds at the folded states. Thus, the specific disulfide bonds play a vital role in controlling the sequences folding into unique 3D structures.

**Table 1.**
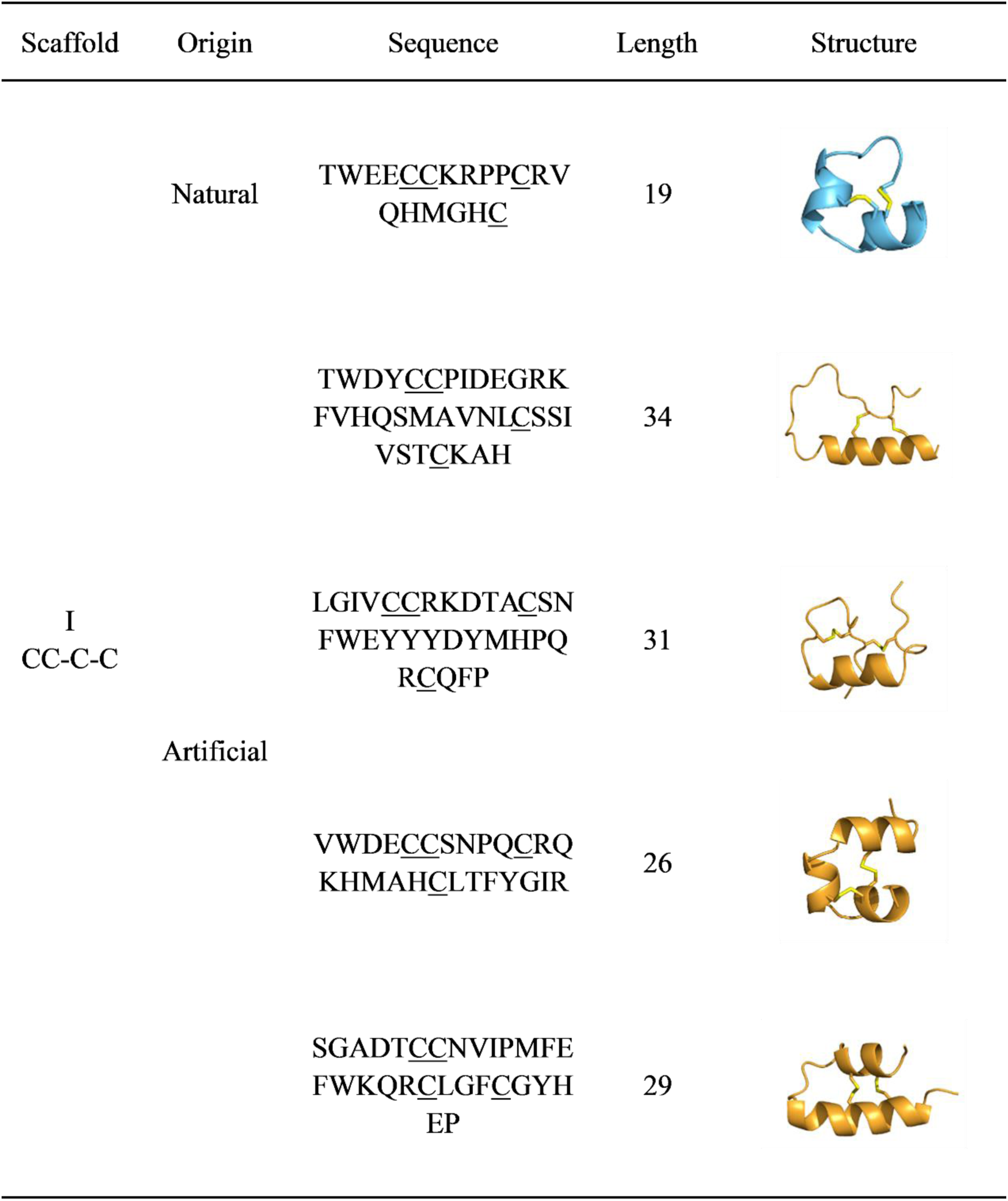
Comparison of the 3D structures between artificial conotoxins and natural conotoxins. (Blue: Natural onotoxins from the ConoServer database; Orange: Artificial conotoxins generated by ConoGen; Yellow: disulfide bond.) Here, the scaffold I (CC-C-C) was shown as an example, the full list can be found in **Supplementary Table 1** (available at https://zenodo.org/records/10679280).

#### Folding stabilities of the different artificial conotoxins

The ability of artificial conotoxin to form a stable spatial structure similar to the natural conotoxin is another crucial aspect in evaluating the quality of ConoGen production. Therefore, we conducted MD simulations on 37 artificial conotoxin structures originating from different cysteine scaffolds. Following the generation of MD trajectories, the stabilities of artificial conotoxins were assessed by calculating the root mean square deviation (RMSD) of Cα atoms, as depicted in Figure 6. After 500 ns of simulation, the majority of MD trajectories reached equilibrium in terms of RMSD. The variation in RMSD for the MD trajectories of 31 artificial conotoxins was below 3 Å, indicating that these artificial conotoxins maintained a relatively stable structure without significant structural changes. However, the RMSD fluctuations in their MD trajectories of the other 6 artificial conotoxins ranged from 4 to 6 Å, suggesting that these conotoxins experienced some degree of change during the simulation but may still have retained a certain overall structure.

**Figure 6.**
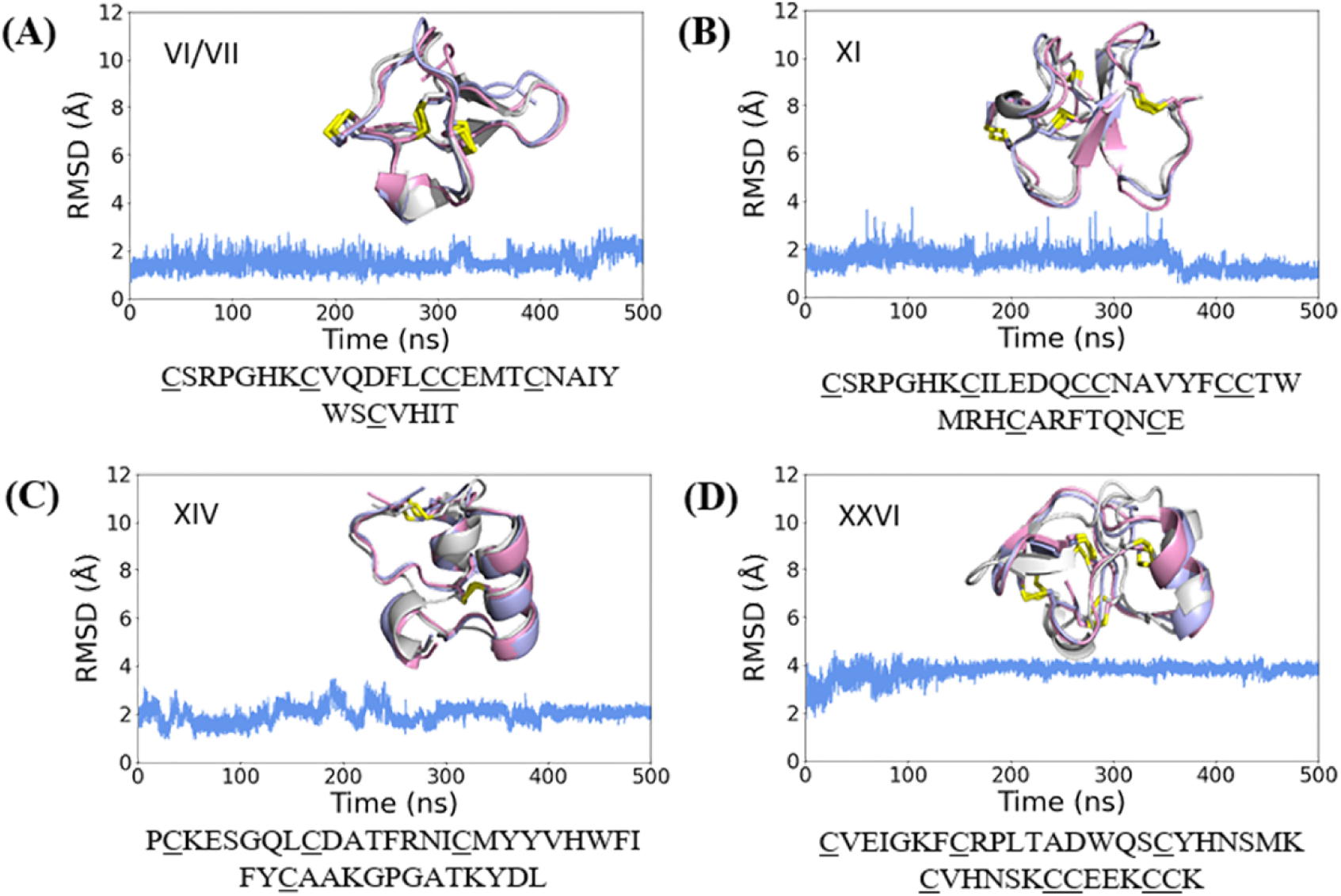
The results of molecular dynamics (MD) simulation of the artificial conotoxin generated by ConoGen. The root mean square deviation (RMSD) of MD trajectory and the alignment of structures from the MD simulation and AlphaFold2 prediction (initial structure) were depicted. (Gray: Initial structure; Pink: Structure extracted at 300 ns during MD simulation; Purple: Structure extracted at 500 ns during MD simulation). Here, the scaffolds **(A)**VI/VII, **(B)** XI, **(C)** XIV, and **(D)** XXVI were shown as examples, the full list can be found in **Supplementary Table 2** (available at https://zenodo.org/records/10679280).

In addition, structures at various time points during the MD simulation were extracted for each simulation trajectory. Further analysis of these artificial conotoxins was conducted by comparing the initial structures before the MD simulation with those obtained during the simulation period. In accordance with Figure 6, the artificial conotoxins with RMSD below 3 Å maintained a relatively stable structure throughout the simulation period. The structure during the simulation closely resembled that of the pre-simulation state. For artificial conotoxins with RMSD fluctuations in the range of 4-6 Å, there were typically long segments without disulfide bonds. The regions lacking disulfide bonds experience structural variations, while the areas with disulfide bonds maintain relative stability throughout the simulation period. Consequently, the substantial fluctuations in RMSD values for these artificial conotoxins can be largely attributed to the presence of long segments lacking disulfide bonds, which lead to increased molecular flexibility during the simulation. In sum, the artificial conotoxins generated by ConoGen not only have the capability to fold into specific spatial conformations, but also exhibit a certain level of stability.

## Conclusions

In this work, we propose ConoDL, a framework for the generation and prediction of artificial conotoxins. ConoDL consists of the conotoxin generation model, ConoGen, and the conotoxin prediction model, ConoPred. ConoGen, inspired by the large language model ProGen, is fine-tuned on a conotoxin dataset based on ProGen. This addresses the challenges posed by the limited conotoxin dataset. ConoPred is constructed based on a neural network architecture using WAE, demonstrating robust predictive capabilities. We can employ ConoPred to preliminarily identify artificial conotoxins, narrowing down the scope for subsequent research and enhancing accuracy. Furthermore, we have demonstrated the feasibility and superiority of ConoDL from both the sequence and spatial perspectives. From the sequence perspective, artificial conotoxins exhibit similarities with natural conotoxins in terms of sequence length range, cysteine framework, and physicochemical properties. Simultaneously, they present a degree of novelty, exemplified by the inclusion of novel cysteine scaffolds. From a spatial perspective, artificial conotoxins can form a specific number of disulfide bonds, folding into a defined structure, and exhibit a certain degree of structural stability. Therefore, ConoDL can address current challenges and expedite exploration within the vast molecular space of conotoxins. Furthermore, ConoDL is poised to be applied in the development of other toxins, making a definite contribution to pharmaceutical research. However, ConoDL still has its limitations. Specifically, Conopred’s predictions are presently limited to the sequence identification stage, lacking the capacity to conduct targeted screening of conotoxins by predicting their pharmacological targets. To overcome this, our future strategy includes developing and implementing a specialized conotoxin target prediction model to more comprehensively explore the vast molecular space of conotoxins.

## Methods

### Dataset preparation

To train ConoGen, a total of 3281 mature conotoxin peptides from the ConoServer database were first downloaded^22^. The most prominent characteristic of these peptides is the presence of a specific number of cysteines residues, which enables them to readily form disulfide bonds, bestowing them with unique 3D conformations^21^. When curating this dataset, we set a criterion to select peptides with lengths ranging from 10 to 70 amino acids. This range encompasses the majority of Conotoxins and is also conducive for the subsequent neural network model training. As a further pre-processing step, the sequences with non-natural amino acids (B, J, O, U, X, and Z) and the ones with lower case letters were considered in this dataset^19^. Furthermore, only peptides containing at least four cysteines were considered, ensuring that the selected peptides have the potential to form a minimum of two stable disulfide bonds^31^. After screening, the dataset used for training ConoGen comprises 2310 conotoxin sequences.

To train ConoPred, we employed the aforementioned 2310 conotoxin sequences as positive samples, and 13,941 protein sequences from Swiss-Prot subset in UniProt database^32^ (https://www.uniprot.org) served as negative samples. These negative samples underwent similar preprocessing to exclude sequences containing any annotations related to conotoxins and sequences containing non-natural amino acids.

### Acquisition of the pre-trained model ProGen

The conotoxin generation model was fine-tuned on the basis of ProGen^15^ using the conotoxin training set. ProGen is a large language model trained using a universal protein sequence dataset, which comprises 2.81 billion non-redundant protein sequences, spanning approximately 19,000 distinct protein families. Herein, pre-trained model (progen2-small) and the associated code was downloaded from Github (https://github.com/salesforce/progen/).

### Fine-tuning ConoGen with conotoxin dataset

In this study, we fine-tuned the pre-training model of ProGen^15^ using the conotoxin dataset. During the fine-tuning stage, the optimization objective is to minimize the negative log-likelihood loss. Specifically, the negative log-likelihood loss for the entire dataset is computed by summing or averaging the losses across all samples. At the same time, the model applied the AdamW optimization algorithm^33^, an adaptive learning rate method that automatically adjusts the learning rate to enhance the stability and speed of the model during the instantaneous descent process. The model was fit for 15 epochs with a learning rate of 0.0001, batch size of 2. All sequences are truncated to a maximum length of 70. Sequences shorter than 70 were padded, and the padding tokens were excluded from the cost function used for training. After the fine-tuning training, the checkpoint with the minimum loss was selected as the final model for conotoxin sampling, named “ConoGen”. Models in this work were trained on a NIVDIA GTX 3080. Programming language: Python. Other requirements: Python3.7, PyTorch, RDKit.

### Generating Artificial Conotoxins Using ConoGen

Next, We utilized ConoGen to generate artificial conotoxins using Top-p sampling. The hyperparameter configuration included “bad_words_ids” set to [(27), (18), (6), (24), (29)] to exclude non-natural amino acids (B, J, O, U, X, and Z). The “Max length” was set to 70, and the repetition penalty adjusted between 1.5 and 1.8. The “P” value ranged from 0.6 to 1.0, and the “T” value ranged from 0.8 to 1.2, both adjusted in increments of 0.1. Each sampling generated 10,000 sequences, totaling 1 million artificial conotoxin sequences.

### Training ConoPred for prediction artificial conotoxin

In order to preliminary identification the artificial conotoxins generated by ConoGen, eliminating less plausible sequences to narrow down the scope of subsequent studies, we trained a conotoxin prediction model, ConoPred. The sequence-based predictive model requires a two-stage training process, which includes training the WAE model and training the classifier. First, the WAE model was fit for 500 epochs with a learning rate of 0.001, batch size of 32. All sequences were truncated to a maximum length of 70. The model with the lowest loss value was selected. The optimal loss value obtained for the WAE model is 1.531. Subsequently, the classifier model was fit for 100 epochs with a learning rate of 0.0001 and a batch size of 32. The model with the highest AUC and AUPR values was selected. The best model achieved an AUC of 1.0 and an AUPR of 0.998 on the training set, as well as an AUC of 1.0 and an AUPR of 0.998 on the test set. The trained WAE model and the classifier model were combined to create a prediction model (named ConoPred) for predicting the probability of conotoxins. Upon inputting sequences into ConoPred, the prediction model will output a probability value for each sequence. This value represents the probability of the sequence being considered a conotoxin sequence, thereby providing a crucial reference for conotoxin selection.

### Evaluation of artificial conotoxins

#### Sequences characteristics and physicochemical properties analysis

The sequence characteristics, including sequence length distribution and cysteine scaffold distribution, as well as physicochemical properties like amino acids distribution, hydrophobicity and net charge, are analyzed and computed using codes developed in-house.

#### Sequence similarity analysis

To analyze the similarity and novelty of artificial conotoxins generated by ConoGen, code was employed to calculate the similarity between natural conotoxins and artificial conotoxins, as well as the self-similarity among the artificial conotoxins. The code used for calculating similarity utilized the pairwise2 module of the Biopython ^20^.

#### 3D structure prediction

ColabFold^34^, a notebook-based implementation of AlphaFold2, was employed to predict the structure of artificially generated sequences by ConoGen. The prediction of the structure was primarily conducted using default parameters, including use_amber (No), template mode (None), msa_mode (MMSeq2 with UniRef and Environmental databases), and num_recycle (3 cycles). The cysteine scaffold types of the artificial conotoxin sequences generated by ConoGen are categorized into 30 classes, with 29 classes conforming to existing cysteine scaffold types, while the remaining class comprises all sequences that do not adhere to existing cysteine scaffold types. One to two sequences were selected for structural prediction from each category, resulting in a total of 34 artificial conotoxin sequences subjected to structural prediction.

#### Molecular dynamics simulation

To understand the structural stability of generated artificial conotoxins, we conducted comprehensive all-atom MD simulations using AMBER16 package. The starting coordinates of artificial conotoxins were obtained from the AlphaFold2 predicted structures. The protein was descripted by AMBER ff14SB^35^. Initially, disulfide bonds were constrained, followed by the addition of an appropriate number of counterions as necessary to ensure the simulation system attained an electrically neutral state. Subsequently, the system was solvated in a cubic box with a side length of 15 Å using the TIP3P water model^35^. Finally, topology and coordinate files were generated and submitted for 500 ns MD simulations. During the simulation, the trajectory was saved every 1 ns, and a time step of 0.002 ps was employed for all simulations. The evolution of the peptides’ Cα atoms root mean square deviation (RMSD) values along the simulations were calculated to evaluate the stabilities of the 3D structures.

## Data availability

The conotoxins data used in this work can be obtained from the ConoServer database (https://conoserver.org/), and peptides serving as negative samples were sourced from the UniProt database (https://www.uniprot.org/). The pre-trained model employed for fine-tuning ConoGen was derived from ProGen (https://zenodo.org/records/7309036). The input files, parameter files, topology files and trajectory analysis scripts used to generate the results in this work, as well as the representative structure of each simulation were provided as supplementary data.

## Code availability

The code of ConoDL and other in-house script for data analysis are open for academic usage and available in GitHub (https://github.com/xueww/ConoDL). Additionally, the supplementary material and model are deposited on Zenodo (https://zenodo.org/records/10679280).

## Notes

The authors declare no competing financial interests.

## Acknowledgements

This work was supported by the Natural Science Foundation of China (21505009), the Natural Science Foundation of Chongqing (2023NSCQ-MSX0140), the Entrepreneurship and Innovation Support Plan for Chinese Overseas Students of Chongqing (cx2020127), the Open Project of Central Nervous System Drug Key Laboratory of Sichuan Province (230012-01SZ).

## Contributions

Weiwei Xue and Zengpeng Li conceived the project. Menghan Guo designed the generative model, performed experiments and analyzed the results. Xuejin Deng contributed to the structural analysis. Ding Luo and Jingyi Yang assist molecular dynamics simulations. Weiwei Xue and Yingjun Chen supervised the project. Menghan Guo developed the first draft of the paper. All authors contributed to writing and improving the paper and approved the submission.

